# Proteome and Secretome Dynamics of Stem Cell-Derived Retinal Pigmented Epithelium in Response to Acute and Chronic ROS

**DOI:** 10.1101/529271

**Authors:** Jesse G. Meyer, Thelma Garcia, Birgit Schilling, Bradford W. Gibson, Deepak A. Lamba

## Abstract

Age-related macular degeneration (AMD) is the leading cause of blindness in developed countries, and is characterized by slow retinal degeneration linked to chronic oxidative stress in the retinal pigmented epithelium (RPE). The exact molecular mechanisms that lead to RPE death and dysfunction in response to chronic reactive oxygen species (ROS) are still unclear. In this work, human stem cell-derived RPE samples were treated with a low dose of paraquat (PQ) for 1 week or 3 weeks to induce chronic reactive oxygen species (ROS) stress. Cells were then harvested and both the intracellular and secreted RPE proteomes were quantified by mass spectrometry. Inside the RPE, chronic ROS caused concerted increase of glycolytic proteins but decreased mitochondrial proteins, as well as decreased extracellular matrix proteins and membrane proteins required for endocytosis. From the secreted proteins, we found that stressed RPE secrete over 1,000 detectable proteins, and the composition of the proteins secreted from RPE changes due to chronic ROS. Notably, secreted APOE is decreased 4-fold due to 3 weeks of chronic ROS stress, and urotensin-II, the strongest known vasoconstrictor, doubles. Further, secreted TGF-beta is increased, and its cognate signaler BMP1 decreased in the secretome. Together, these alterations of the RPE proteome and protein secretome paint a detailed molecular picture of the retinal stress response in space and time.

## Introduction

AMD is the leading cause of blindness in people over age 50, and represents an area of significant unmet clinical need. AMD is characterized by retinal degeneration in the center of the retina, the macula. Three tissues comprise a minimally functional unit of the retina, RPE is the epithelial layer between the light-sensitive PRs and vasculature (choroid). RPE is especially important among this triplet because it forms the outer blood-retinal barrier due to the tight-junctions between the cells. RPE is also the main support layer for the PRs and some RPE functions relevant to photoreceptor survival and function include: (i) receipt of nutrients from vasculature and transport of nutrients to PRs^1^, (ii) phagocytosis of shed photoreceptor outer segments, and (iii) secretion of signals including growth factors and cytokines^2^. The functional disruption and atrophy of the RPE is a key factor in the progression of degenerative conditions in the retina, leading to the death of other cell types in the retina, including the light-sensitive rod and cone PRs, resulting in significant vision loss. Enabling study of retinal diseases, RPE can be generated from induced pluripotent stem cells (iPSCs)^3,4,^ using small molecules and growth factors that mimic developmental cues.

Progression of AMD is associated with chronic ROS especially in the RPE layer^5^. ROS in retina is produced mostly due to high rates of metabolism in RPE^6^ and the resulting electron leakage from mitochondria^7^. Notably, a large amount of oxidative molecules may result from phagocytosis of PR outer segments^8^. Overall, the main site of oxidative injury appears to be the mitochondria, and pathological studies suggest that RPE damage is an early event in AMD^9^, which justifies the need to understand the oxidative stress response mechanisms in RPE. Chronic oxidative damage in RPE may increase the expression of proteins (see ^10^) and probably causes apoptosis. Stem cell-derived RPE was previously used as a model system with chronic, low-level oxidative stress from paraquat (PQ) to study the NRF2-mediated transcriptional responses^11^. PQ is well tolerated by RPE^12^ and is a suitable mimic for pathological oxidative stress that occurs in AMD, which is thought to be mostly generated in mitochondria^13^. Although the connection between oxidative stress and RPE dysfunction is clear, the exact molecular changes the mediate dysfunction remain poorly defined.

Altered protein secretion and the related changes in extracellular matrix organization (ECM) are also known to be involved in AMD, especially loss of barrier function and vascular invasion^14^. Several reports describe RPE secretion of various proteins, including ARMS2^15^, which is secreted and interacts with other proteins that are mutated in AMD, such as fibulin-6, although the explicit function of secreted ARMS2 was not determined. Other studies using RPE cells have looked at specific protein secretions, such as apolipoprotein E (APOE)^16,17^. Finally, the dry form of AMD is associated with deposits rich in lipids and proteins called drusen. The protein components of drusen were found to include several complement proteins and APOE^18^. One recent study also reported use of iPSC-derived RPE to study altered transcription and protein secretion^19^. What is not well understood is how the RPE secretome changes over time during chronic oxidative stress to communicate its state to adjacent PR and vasculature.

Although much is known about the pathology of AMD, we still do not understand the exact molecular mechanisms that lead from chronic stress to tissue dysfunction and the eventual pathology described above. In such cases, unbiased, system-wide measurements of molecular remodeling in response to stress can help provide new ideas for mechanistic follow-up experiments. Mass spectrometry-based proteomics is one such method^20^. Recent advancements, particularly using data-independent acquisition (DIA)^21^ and faster hardware, have enabled quantification of over 5,000 proteins per hour^22–25^. This strategy provides more complete data across replicate samples with excellent quantitative accuracy, thereby enabling more comprehensive discovery of biological pathways.

Here we used stem cell-derived RPE tissue as a model system and performed unbiased DIA mass spectrometry-based proteomics to understand the intracellular mechanisms that mediate RPE ROS tolerance, and how those changes are communicated to surrounding cells through secreted proteins (the secretome). We collected data from the intracellular proteome and the secretome after 1 or 3 weeks of treatment to simulate acute and chronic stresses. Further, to understand how RPE adapts from acute to chronic ROS, we compare the proteome and secretome after 3 weeks of stress with that after one week. Our results show that low-level PQ treatment of RPE for 1 or 3 weeks causes significant alterations of metabolic proteins, suggestive of a shift in RPE from lipid to sugar metabolism. We also find a number of protein changes related to cellular organization and translation. From the secretome data, we find that acute and chronic ROS significantly alter proteins in the pathways of ECM-receptor interaction, focal adhesion, and complement and coagulation cascades. Remarkably, we also found significant decreases in APOE, amyloid-precursor-like protein, and the strongest known vasoconstrictor urotensin-2. Together, our results paint a detailed picture of the human RPE’s proteomic landscape in response to acute and chronic ROS stress over space and time that point to possible mechanisms of cellular damage in AMD.

## Results

### Workflow and General Overview

An overview of the experimental strategy is shown in Figure 1A. RPE derived from hESC were treated with PQ for 1 or 3 weeks, and the conditioned media containing secreted proteins was collected to quantify changes in the composition of secreted proteins (secretome, SEC). Parallel experiments were carried out where RPE cells were harvested in order to quantify the intracellular proteome changes. We collected DIA data from each sample and used Spectronaut^26^ with the pan-human library^23^ to identify and quantify proteins. Using this strategy, we identified a total of 3,257 proteins from the intracellular proteome samples, and a total of 1,319 proteins from the secretome samples. Among those proteins, 1,054 proteins were identified in both groups (Figure 1B). We assessed the quality of our proteome quantification by comparing the coefficients of variation for all identified protein groups within each treatment group (Figure 1C and 1D). Based on the proportions of each group that had CV < 20% or <10%, the quality of the data from the intracellular proteome quantification was greater than the quality of the secretome. This difference in variability among the RPE and SEC experiments is reflected in the number of protein changes we were able to detect from each experiment (Figure 1E, **Supplementary Table 1**). For both the RPE and SEC experiments, 1-week or 3-week PQ treatments were compared to controls. In addition, 3-week treatment results were also compared to the 1-week treatment data in order to determine which proteins might be involved in the transition from acute and chronic ROS adaptation.

**Figure 1:**
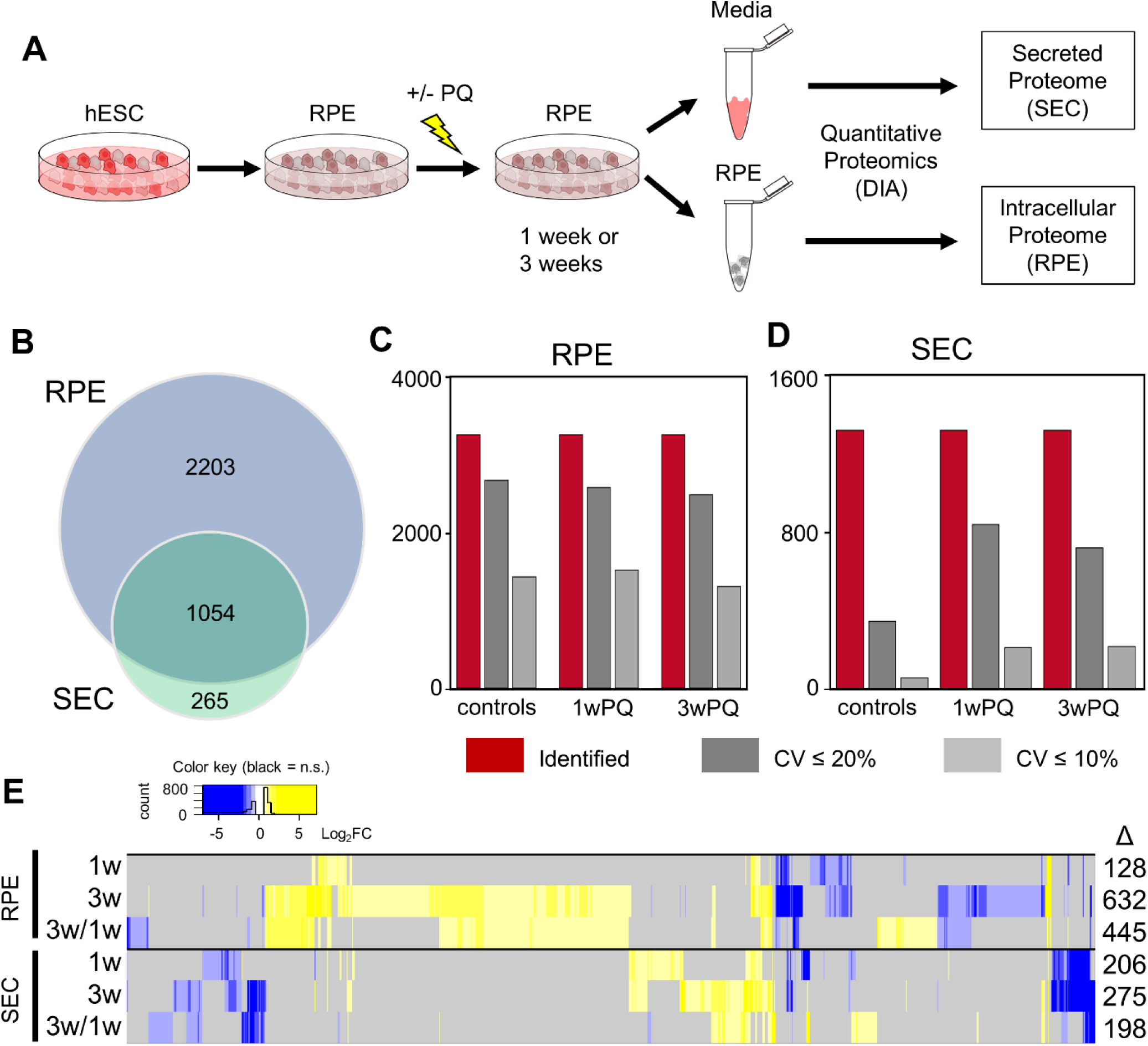
Proteomics study workflow and results overview. **(A)** Human embryonic stem cells (hESC) were differentiated into retinal pigmented epithelium (RPE) cells, allowed to mature, and then treated with paraquat (PQ) or vehicle control for 1 week or 3 weeks. After treatment, serum-free media was collected to measure the protein secretome (SEC), and cells were harvested to measure the intracellular proteome (RPE). (**B)** Venn diagram showing the number and overlap of proteins identified in the two datasets. (**C)** Subsets of identified and quantified protein groups from each RPE treatment condition showing the total in maroon, those with less than 20% coefficient of variation (CV) in dark grey, and those with <10% CV in light grey. (**D)** The same as (**C)** for SEC data. (**E)** Heatmap showing overview of statistically significant changes (q-value< 0.01, fold-change > 50%) found in each of the comparisons.

### Intracellular Proteome Changes

To understand how RPE changes in response to acute or chronic ROS, we first looked more closely at the intracellular proteomic changes. After 1 week of PQ treatment, only 128 proteins were altered in abundance. KEGG pathway enrichment analysis of those proteins revealed only a handful of pathways: the citric acid cycle, glycolysis, and a cluster of pathways related to oxidative phosphorylation (OxPhos, Figure 2A). All of these pathways are related to cellular energy production. Notably, the direction of protein changes in these pathways clearly suggests a decrease in mitochondrial metabolism and an increase in glycolysis. After 3 weeks of ROS stress in RPE, the changes were much more pronounced. Over 600 proteins showed expression changes, and could be mapped to 20 KEGG pathways (Figure 2B). Again, most of these pathways were associated with energy production, except for aminoacyl-tRNA biosynthesis, which was found to mostly increase and is related to protein synthesis. Many additional pathways related to amino acid and fatty acid metabolism were also found, in addition to an increase of proteins needed for glycolysis. However, after the 3-week PQ treatment the changes to the cluster of OxPhos pathways were more pronounced and complex likely due to activation of cellular compensatory mechanisms to chronic oxidative stress. To better understand how acute and chronic ROS from PQ treatment were influencing the mitochondrial electron transport chain (ETC), we generated a model of ETC complexes and colored their subunits according to whether they were found to be increased or decreased (Figure 2C-D). Remarkably, acute ROS caused a concerted decrease in subunits of ETC complex I, as well as a decrease of complex II subunits SDHA and SDHB (Figure 2C). This result supports reports that complex I is the major site of ROS production by paraquat^27^. Chronic ROS also caused a nearly concerted decrease in ETC complex I subunits; however, concerted increases in subunits of complexes III, IV, and V were now also apparent (Figure 2D). This data suggests that cells compensate for the reduction in complex I activity with an increased expression of downstream subunits.

**Figure 2:**
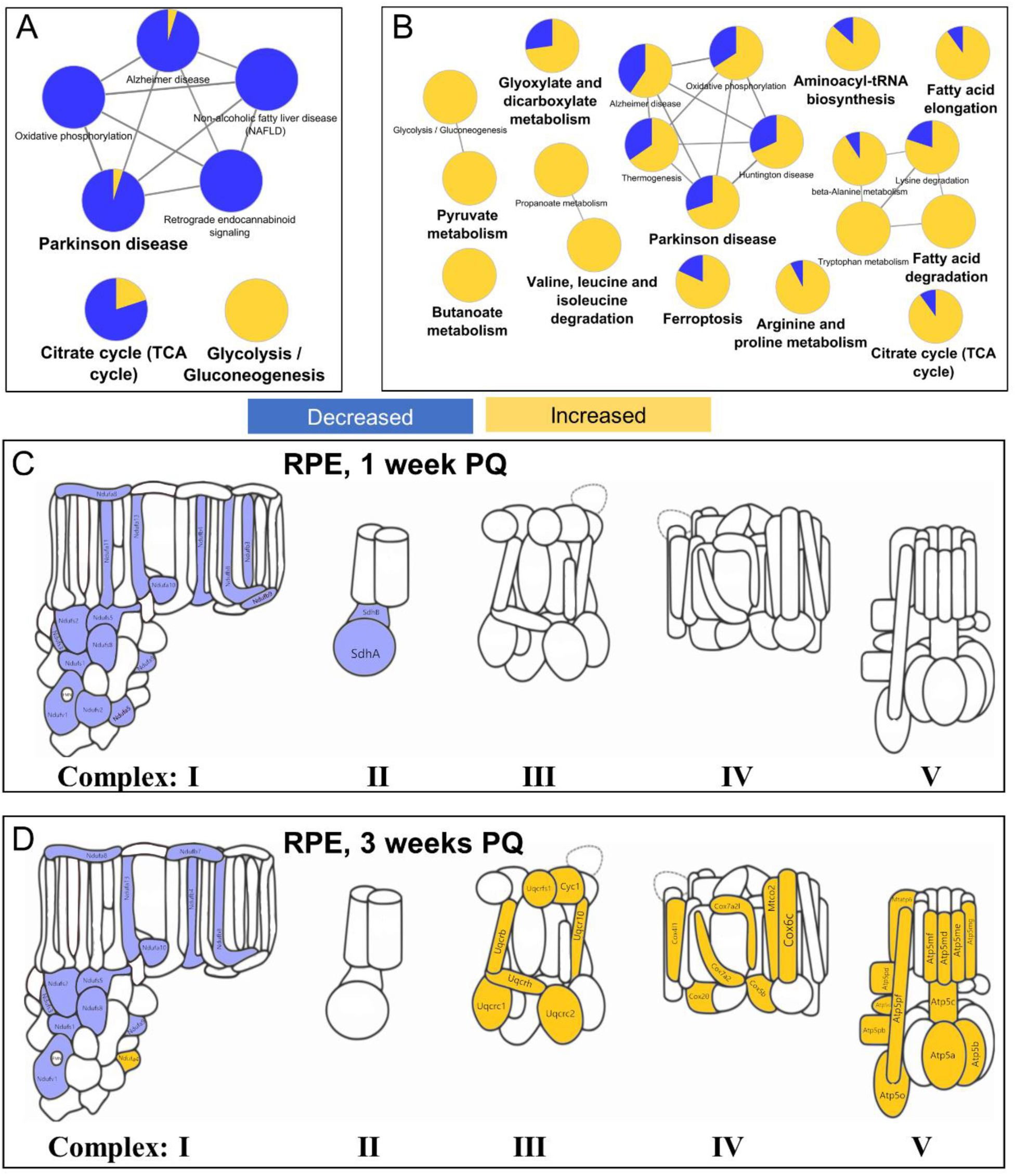
RPE intracellular proteome changes resulting from chronic ROS. KEGG pathway analysis showing enriched pathways (p-value < 0.001) of the intracellular RPE proteome changes after (A) 1 week or (B) 3 weeks of treatment with PQ. Subunits of the electron transport chain showing statistically significant changes after (C) 1 week or (D) 3 weeks of treatment with PQ.

The most- and least-changed intracellular proteins after 3 weeks of ROS stress do not appear in the pathway enrichment analysis. The most increased proteins were GDF15, PLIN2, and SQSTM1 (**Figure S1A**). Known NRF2-regulated proteins NQO1, SRXN1, and HMOX1 are also found in the top 6 increased proteins^11,28^. The most decreased proteins were COL6A1, LRP2, PCBP4, CD74, GLUL, and the complement cascade inhibitor, SERPING1 (**Figure S1B**).

### Intracellular Adaptation from Acute and Chronic ROS

Because we collected proteomic data from both 1-week and 3-week ROS treatments, we can also uniquely assess how ROS exposure evolves from acute to chronic response by comparing the protein quantities directly between these two time points (3w/1w). In order to understand only the adaptation of the ROS response from acute to chronic we excluded any proteins that changed in the 1w/controls or 3w/controls comparison, leaving 92 protein changes. KEGG pathway enrichment analysis using this subset of changes found Glyoxylate and dicarboxylate metabolism and Propanoate metabolism pathways (enrichment p-values < 5e-4, Figure 3A). Only 5 proteins (CAT, GLDC, MUT, ABAT ACSS3) were found in these two pathways, all of which were increased. To explore if there are other differences, we also checked for enriched wikipathways (Figure 3B), and found the ETC pathway (enrichment p-value = 4e-7), including SCO1 and SURF1 which assemble cytochrome C oxidase. Finally, the set of altered proteins were checked for enrichment of gene ontology (GO) molecular functions and biological processes (Figure 3C). Notably, several proteins involved in mitochondrial membrane organization and protein import, as well as several mitochondrial ribosomal proteins translation were increased. Among these proteins, alpha-synuclein and NDUFA7 were the only downregulated members. Proteins with peroxidase activity (Catalase, MGST2 and GPX8) were also upregulated.

**Figure 3:**
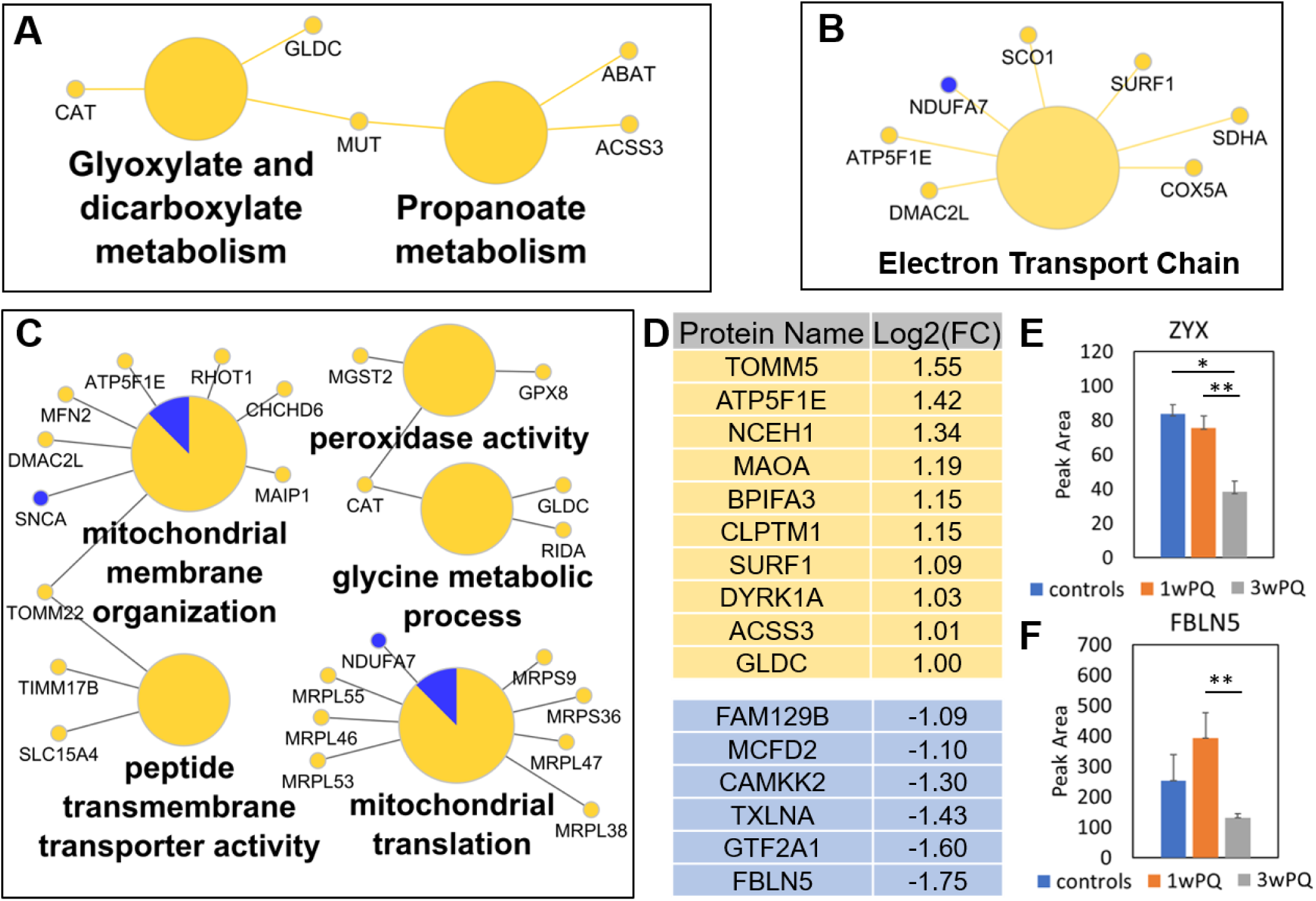
Temporal dynamics of RPE intracellular proteome from 1 week to 3 weeks of chronic ROS. Proteins that changed only in the 3-weeks / 1-week comparison found enriched in (A) KEGG pathways, (B) Wikipathways, or (C) Gene Ontology Molecular Function and Biological Processes. All pathway enrichment analysis used a minimum p-value of 0.001. (D) List of proteins that increased or decreased at least 2-fold. (E) Quantification of Zyxin protein from the average of the SPGAPGPLTLK^2+^, QNVAVNELCGR^2+^, and FGPVVAPK^2+^. (F) Quantification of Fibulin-5 protein from the peptide DQPFTILYR^2+^. Error bars show the standard error. Significant changes defined as at least 1.5-fold and q-value<0.05 (*), q-value < 0.01 (**), and q-value<0.001 (***).

In the list of altered proteins there were many interesting changes that were not part of these concerted pathway changes. Of these, two categories jump out: calcium signaling and structural protein changes. Among the significantly decreased proteins were structural proteins Zyxin (Zyx, UniProt: Q15942, Figure 3E) and fibulin 5 (FBLN5, UniProt: Q9UBX5, Figure 3F). The latter was the most downregulated protein. FBLN5 is especially interesting in the context of AMD because it localizes to Bruch’s membrane in the eye^29^ and mutations in FBLN5 are known to be associated with AMD^30,31^. Three of the most decreased proteins are involved in calcium signaling, Multiple coagulation factor deficiency protein 2 (binds to calcium ions)^32^, alpha-taxilin (may play a role in calcium-dependent exocytosis)^33^, and calcium/calmodulin-dependent protein kinase kinase 2. Related to calcium signaling, a member of the calcium-activated chloride channels, Anoctamin-10^34^ (UniProt: Q9NW15), was increased almost 2-fold (**Supplementary Table 1)**. Finally, in the lists of the most increased and decreased proteins (at least 2-fold change from 3-weeks /1 week), several proteins have yet unknown function in the retina, such as CLPTM1, BPIFA3, MAOA, and FAM129B (Figure 3D).

### Secretome Remodeling

Next, we examined the remodeling of secreted proteins. Protein changes representative of several KEGG pathways changed after 1 week of PQ treatment, including “complement and coagulation cascades”, “protein digestion and absorption”, and a cluster related to ‘ECM-receptor interaction” and “Focal adhesion” (Figure 4A). After 3 weeks of PQ treatment, proteins in the glycolysis and lysosome pathways were also found to increase in the extracellular medium (Figure 4B). In common between the 1-week and 3-week changes were the cell and ECM structure-related pathways of “ECM-receptor interaction”, “focal adhesion”, “amoebiasis”, as well as the “complement and coagulation cascades”. Figure 4C shows these common pathways plotted with the protein changes that make up and connect these pathways. Related to the structural protein pathways, we see decreases in several intracellular and extracellular organization proteins, including actin, actinin, thrombospondins 1 and 2, fibronectin, and collagens. The Laminin changes were mixed, with subunits alpha-5 and beta-2 increased, but alpha-4 and beta-1 were decreased. Within the complement cascade several factors were decreased, including: CFI, CFB, C3, C4A, and C4B. Overall, the data indicates significant deregulation of the protein machinery for the complement, coagulation, and organizational systems after chronic ROS stress.

**Figure 4:**
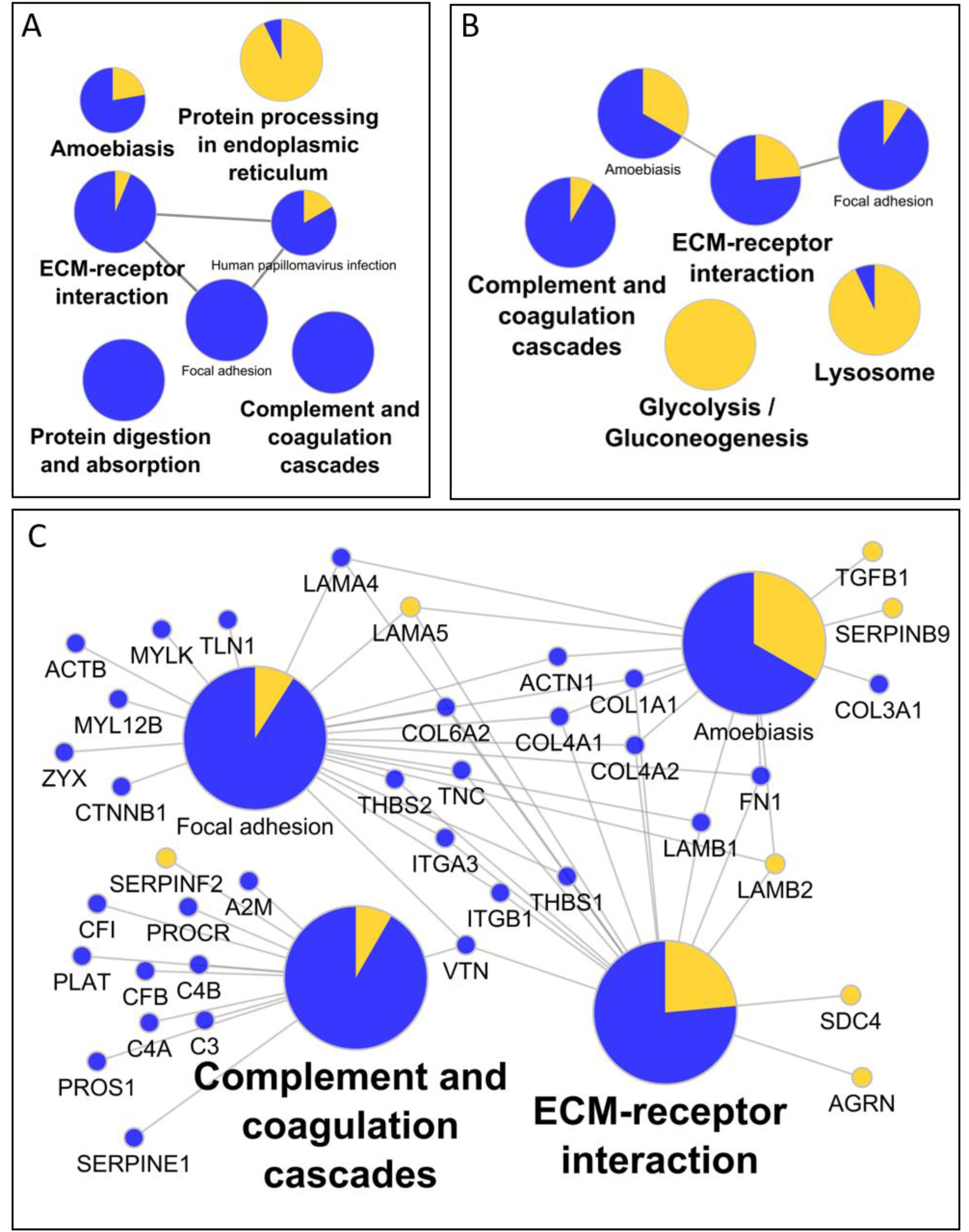
Functional characteristics of secreted protein changes. KEGG pathways of proteins altered after (A) 1 weeks or (B) 3 weeks of PQ treatment. (C) Expansion of proteins related to ‘ECM-receptor interaction’ and ‘Complement and Coagulation Cascades’ that change after 3 weeks of PQ treatment.

There are several notable protein changes among the secretome quantification that did not appear in the pathway analysis. Since we noticed an increase in TGF-β1 (Figure 4C, **Figure S2Ai**) we looked in the list for related extracellular signaling proteins, and found that BMP1 decreased three- to four-fold with acute and chronic treatment (**Figure S2Aii**). TGF-β2 was decreased more subtly around 30% after 1 or 3 weeks of stress (**Figure S2Aiii**), which was outside the cutoff used for the main analysis but can be found in **Supplemental Table 4**. Also altered was apolipoprotein E (APOE), which has been recently linked to AMD wherein the APOE2 allele increases risk, while APOE4 is protective, in contrast to Alzheimer’s disease^35,36^. We observed that APOE was down over 2-fold after 1 week of stress, and nearly 4-fold after 3 weeks of stress (**Figure S2Bi**). Another related protein strongly associated with high myopia, amyloid β-like protein 2 (APLP2), was significantly increased after 3 weeks of stress (**Figure S2Bii**). Finally, given that development of AMD is associated with both loss and gain of vasculature, it is notable that urotensin-II (UTS2), the strongest known vasoconstrictor, was more than tripled after 1 or 3 weeks of PQ induced stress (**Figure S2Biii**).

Given the apparent ECM remodeling we also checked for any changes in secreted proteases. We found a large number of proteases and protease-related proteins altered in the secretome data. We did identify two matrix metalloproteases (MMPs), MMP2 and MMP15, but only MMP2, a type IV collagenase, was significantly decreased (**Figure S2Ci**). One of the most upregulated proteases was ADAM9, a known ‘sheddase’ protease that cleaves extracellular regions of membrane proteins^37^ and is known to regulate pathologic angiogenesis in the retina^38^ (**Figure S2Cii**). Finally, a serine carboxypeptidase, Retinoid-inducible serine carboxypeptidase (SCPEP1), was increased over 50% after 3 weeks of PQ stress (**Figure S2Ciii**).

### Acute-to-Chronic adaptation of the Secretome

As described for the intracellular proteome data, we can understand the adaptive response in the Secretome using the proteins that only change from 1 week to 3 weeks of stress. This filtering leaves only 29 increased and 34 decreased proteins. Reactome pathway analysis of these altered proteins returned many altered candidate pathways centered around the decreased levels of tubulins (Figure 5A). Overall, there are many less concerted changes in this data subset than the others. However, there are more interesting individual changes. Amyloid precursor protein (APP) was increased between one and three weeks of stress (Figure 5B). The most increased protein in the adaptation of the Secretome was tetranectin (almost 14-fold increase), which is a potential marker of several diseases and may play a role in exocytosis^39–41^. Retinol-binding protein 4 (RBP4) was also increased over 2-fold (Figure 5C). RBP4 plays an important role in facilitating transportation and cellular uptake of retinol ^42^.

**Figure 5:**
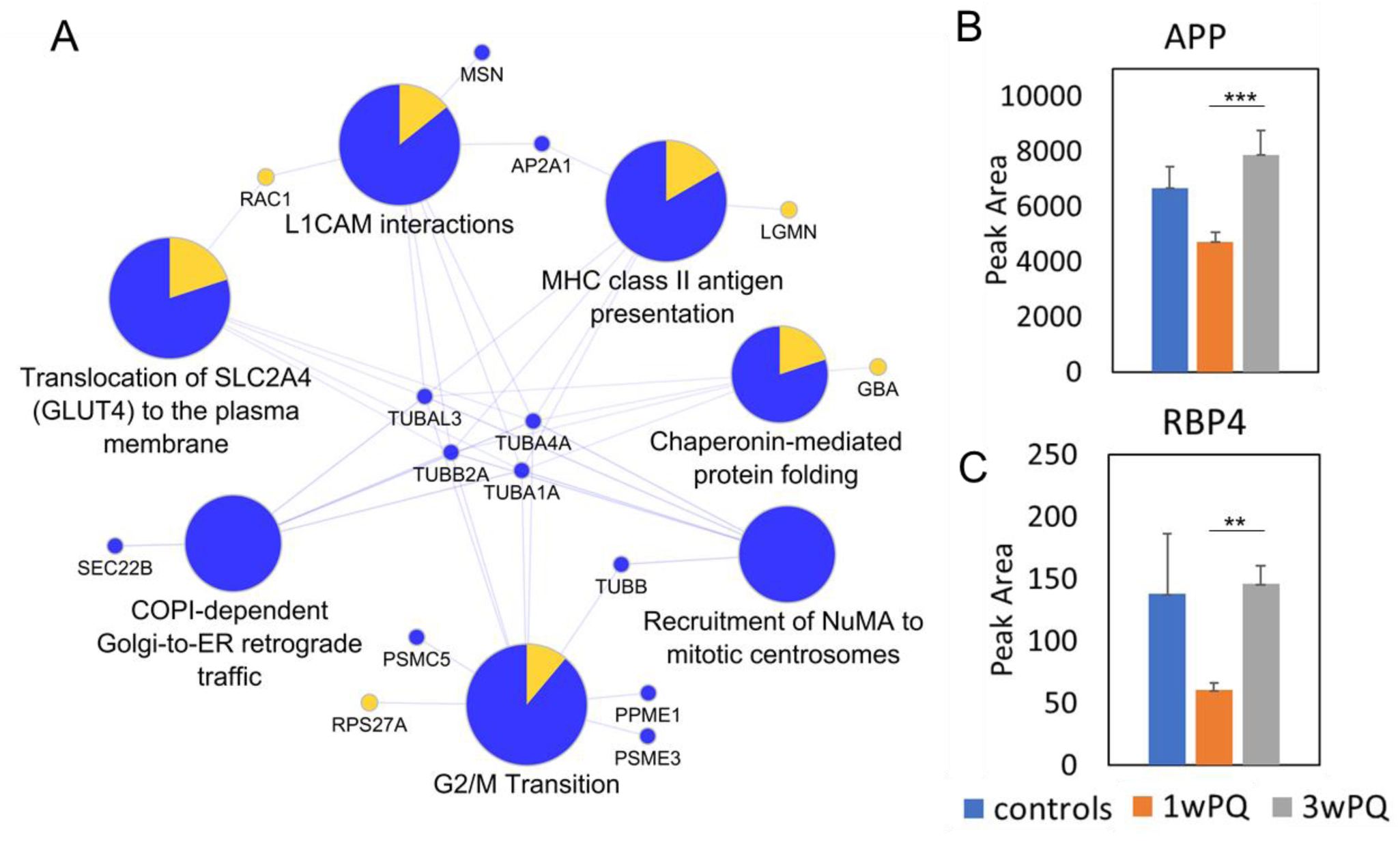
Acute-to-chronic adaptation in secreted protein profiles. (A) Subset of the Reactome pathways enriched in only the Secretome proteins that change from 1 week to 3 weeks of stress (pathway enrichment corrected p-values <0.01). (B) Quantification of Amyloid precursor protein (APP). (C) Quantification of Retinol-binding protein 4 (RBP4). Error bars show the standard error. Significant changes defined as at least 1.5-fold and q-value<0.05 (*), q-value < 0.01 (**), and q-value<0.001 (***).

## Discussion

The results presented here indicate that oxidative stress induces remodeling inside RPE indicative of a metabolic shift from respiration toward glycolysis. A shift to glycolysis has been observed in other systems to reduce oxidative damage under high energy demand^43^. We also found intracellular proteome changes suggestive of ECM and cellular organization changes. Some intracellular protein changes have less-clear implications and require follow-up experiments. For example, CD74, a major histocompatibility class II antigen that acts as a transcription regulator linked to cell survival^44^ was decreased. GDF15, a distant member of the BMP subfamily and target of p53^45^, was also highly upregulated after chronic ROS, although its function in RPE has not been studied.

Acute and chronic oxidative stress also induce significant remodeling of the secreted proteome or secretome. Notably, there were many altered proteins in the complement system which is known to be affected in AMD^16,46,47^. We also observe many changes in proteins related to ECM, focal adhesion, and altered TGFβ signaling^48^, which together may regulate neovascularization. Most interestingly, we find a number of Alzheimer’s-related proteins change in the secretome of oxidatively stressed RPE, including APP, APLP2, and APOE. The different isoforms of APOE have been associated with altered risk of AMD^36,49,50^, and APOE was previously demonstrated to be secreted from RPE and regulate lipid uptake^17^.

It is also interesting to consider the proteins we identified that did not change, such as HTRA1. Single-nucleotide polymorphisms (SNPs) near the ARMS2/HTRA1 genes are associated with altered risk of developing AMD^51–53^. We detected HTRA1 protein both inside RPE and in the secretome. Intracellularly, HTRA1 was increased between 1 and 3 weeks of stress but did not meet our statistical criteria for defining a protein change (39% increase, q-value = 0.016). Extracellularly, HTRA1 was increased 65% but the q-value was only 0.047. We also identified HTRA2 from the intracellular proteome data, and HTRA2 was significantly increased between the chronic stress and control groups according to our cutoffs (54% increase, q-value < 0.001).

Our study uniquely addresses both the temporal and spatial reorganization of the RPE proteome. We expect this data will serve as an important resource for future mechanistic and therapeutic studies. For example, the data suggest new proteins that may mediate the observed retinal disfunction in AMD in tissues adjacent to RPE, the photoreceptors and vasculature (Figure 6). Increased TGFβ1 and decreased BMP1 expression may signal to the photoreceptors to dedifferentiate, or to resident macrophages to influence their pro- or anti-inflammatory state. The decreased APOE secretion may decrease the RPE’s ability to uptake lipids and possibly the shed photoreceptor outer segments. Increased secretion of proteases ADAM9 and SCPEP1 might breakdown the basement membrane, and increased urotensin-II might cause choroidal vasculature to constrict and limit the oxygen delivery to the region. All of these ideas are opportunities for follow-up research uniquely enabled by this study. Taken together, this intracellular and secretome remodeling of RPE cells shown by proteomics provides a highly granular view of the acute and chronic oxidative stress responses.

**Figure 6:**
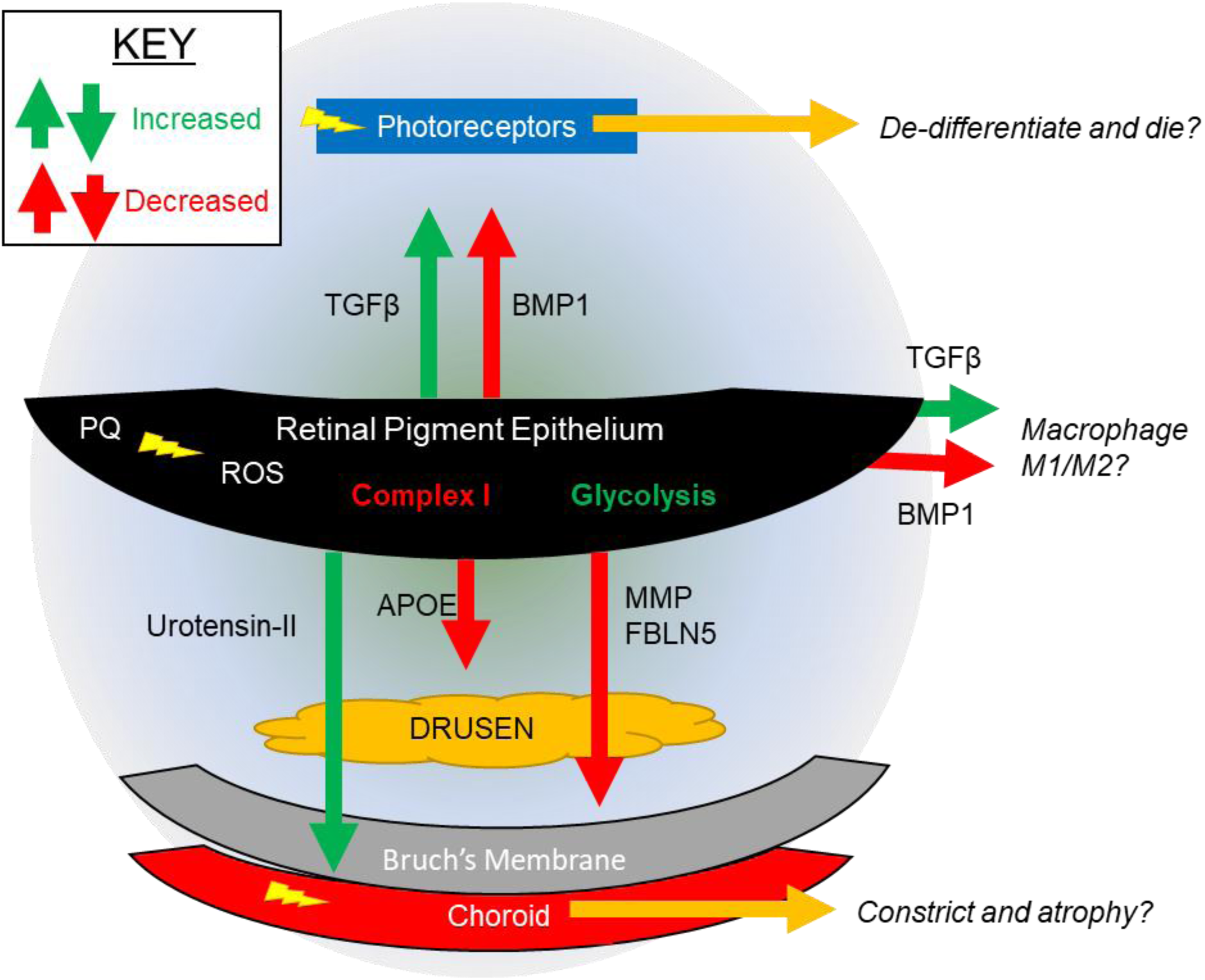
Proposed Spatial Model of RPE Response to Chronic ROS.

## Acknowledgements

JGM was supported by an NIH training grant (T32 AG000266). Work in Dr. Lamba’s lab is supported by NIH grant EY025779. The authors acknowledge support from the NIH shared instrumentation grant for the TripleTOF system at the Buck Institute (1 S10 OD016281).

## METHODS

### Chemicals and reagents

LC-MS grade acetonitrile and water were manufactured by Burdick and Jackson and purchased from Thermo-Fisher. Urea and the BCA assay kit (Pierce) were also purchased from Thermo-Fisher. LC-MS grade formic acid and 1 M triethanolamine bicarbonate (TEAB), pH 8.5, paraquat (PQ) and tris base were purchased from Sigma-Aldrich. Sequencing-grade modified trypsin was purchased from Promega.

### Generation of RPE from human embryonic stem cells

All stem cell work was approved by the Buck Institute SCRO committee. Human ESCs (WA-01; National Institutes of Health registry #0043) were maintained in Essential 8 medium (Gibco, Grand Island, NY, USA) and 1% penicillin-streptomycin-amphotericin B solution (Lonza, Walkersville, MD, USA). Cells were grown on Matrigel (BD Biosciences, St. Paul, MN, USA)-coated plates and serially passaged using 0.5 mM EDTA solution. Genotyping revealed that cells did not have any key AMD single nucleotide polymorphism (SNP)^54,55^. Human ESCs were differentiated to retinal lineage using our previously published protocol^4,56–58^. The RPE regions were manually picked and expanded.

The RPE cells generated from hESCs were cultured in α-MEM medium (Life Technologies, Grand Island, NY, USA) containing 1% fetal bovine serum (Atlanta Biologicals, Flowery Branch, GA, USA), L-glutamine (VWR, Radnor, PA, USA), taurine (Sigma-Aldrich Corp., St. Louis, MO, USA), hydrocortisone (Sigma-Aldrich Corp.), and tri-iodo-thyronine (Sigma-Aldrich Corp.) on Matrigel-coated plates or filter membranes (VWR)^59^. Cells were subcultured using Accutase (Gibco) in the presence of thiazovivin (1 lM), a Rho-associated protein kinase pathway inhibitor that allows passaging RPE cells for over eight passages^60^. The RPE cells were in culture for up to seven passages without any appreciable loss in their ability to mature into polarized RPE cells. The cells were allowed to grow to 100% confluence and used for experiments following maturation.

### Cell Treatments and Collection

RPE cells were treated as described previously^11^. Briefly, RPE cells were seeded in 12-well dishes and allowed to mature for up to 3 weeks following final passage. Fully confluent plates of cells showing typical cobblestone morphology and presence of pigmentation were treated every other day with 160 μM Paraquat (PQ) diluted in RPE media described above for 1 week or 3 weeks. In parallel on the same multi-well dish, control RPE were grown with the same media lacking PQ. After treatment, the cells were washed once with PBS and then trypsinized. Media containing KSR was then added to quench the trypsin activity after 5 minutes, and cells were concentrated by centrifugation. The media was aspirated and the cells were washed once with PBS before being frozen at -80°C until further processing.

### Sample Preparation for Intracellular Proteome Measurement

RPE cell pellets were lysed and proteins were simultaneously precipitated according to the MPLEx protocol ^61^. To each frozen cell pellet, 100 μL of PBS was added, and then 5 volumes of cold (−20°C) chloroform-methanol (2:1 [vol/vol]) was added to each pellet and cells were vortexed until they thawed and dissolved in the solvent. Samples were incubated on ice for 5 minutes, vortexed in the cold room for 5 minutes, and then centrifuged at 15,700 relative centrifugal force (RCF) for 10 minutes at 4°C, which helped clarify a biphasic solution with proteins at the interphase. The top aqueous layer was removed taking care to avoid the protein disc at the interphase, and then 500 μL of cold (−20°C) methanol was added resulting in a monophasic solution. Samples were then vortexed briefly and centrifuged at 15,700 RCF for 10 minutes at 4°C to pellet proteins. The supernatant was removed and the protein pellet was dissolved again with 1 mL of cold (−20°C) methanol by vortexing to wash away any remaining lipids, and proteins were pelleted again by centrifugation at 15,700 RCF for 10 minutes in the cold room. The methanol wash was removed, and the protein pellet was dried briefly in a vacuum centrifuge. The dried protein pellet was then dissolved in 8M Urea and 100 mM TEAB containing protease inhibitor cocktail, and protein concentration was quantified using the BCA assay. Protein disulfides were reduced by adding 4.5 mM DTT (final concentration) and heating to 37 degrees for 30 minutes, and then samples were cooled to room temperature before adding 10 mM iodoacetamide (final concentration) to alkylate free thiols. Samples were then diluted 4-fold with 100 mM TEAB to reduce the Urea concentration, and enzymatic protein hydrolysis was initiated by the addition of trypsin (Sequencing Grade Modified Trypsin, Frozen, Promega) at a ratio of 1:50, trypsin: substrate weight.

### Sample Preparation for Secreted Proteome Measurement

RPE cells were stressed as described above for 1 week or 3 weeks, and then cells were washed three times with 1x PBS before adding 2 mL of serum-free RPE media. Cells were cultured for 24 hours and then the media was collected and frozen at -80°C. Media containing secreted proteome was then concentrated using a centrifugal ultrafiltration membrane (Amicon Ultra-15, MWCO 30 kDa) by centrifugation at 7,500 X gravity for 10 minutes at 4°C. To denature proteins the buffer was exchanged three times by adding 1 mL of 2M Urea in 50 mM Tris, pH 8.0 and centrifuging. The hold-up volume of the buffer-exchanged protein was then removed from the ultrafilter and protein was quantified using the BCA assay. Protein disulfides were then reduced by adding 4.5 mM DTT (final concentration) and heating to 37 degrees for 30 minutes, and then samples were cooled to room temperature before adding 10 mM iodoacetamide (final concentration) to alkylate free thiols. Enzymatic protein hydrolysis was initiated by the addition of trypsin (Sequencing Grade Modified Trypsin, Frozen, Promega) at a ratio of 1:50, trypsin: substrate weight.

### Mass Spectrometry Data Collection and Analysis

Peptides were analyzed by nanoflow liquid chromatography – tandem mass spectrometry analysis (nanoLC-MS/MS) on a Sciex 5600 TripleTOF mass spectrometer using data-independent acquisition (DIA) using variable-width precursor isolation windows as described previously^62^. Peptide separation for DIA data collection was performed using a linear gradient of 5% mobile phase B (98% Acetonitrile, 0.05% formic acid, 1.95% water) to 40% B over 200 minutes with a flow rate of 300 nL/min.

### Data Analysis

Proteins were identified and quantified from DIA-MS data using Spectronaut^26^ and the pan-human spectral library^23^. Spectronaut settings were all defaults except that quantification filtering was set to Q-value percentile = 25% for intracellular proteome, and Q-value percentile = 20% for secreted proteome. Statistically-significant protein changes (defined as q<0.01 and at least 50% change) were assessed for enriched pathways and visualized using the ClueGO plugin of Cytoscape^63^. Additional analysis was carried out using custom scripts in R^64^.

### Sample Groups and Statistical Comparisons

Data from intracellular proteome was collected from three wells of treated cells and two wells of control cells per time point, resulting in a total of 10 samples for intracellular proteome. To ensure that we only detect changes that result from acute or chronic ROS, we compared the treated samples in triplicate with the four control samples, two from each treatment timepoint. Data from the protein secretome was collected from four replicates of treated cells and four replicates of control cells at each timepoint. As described for the intracellular proteomics comparison, samples from each treatment timepoint were compared with a pool of all 8 control samples.

### Data Availability

All raw mass spectrometry data files, tables of quantification, and protein peak areas are available from massive.ucsd.edu under dataset identifier MSV000083127 (password: intrpesec).

### Supplemental Tables

All supplemental tables are also available with the raw mass spectrometry data on Massive or from Zenodo: http://doi.org/10.5281/zenodo.1482734. **Supplemental table 1** gives the list of protein log2(fold changes) filtered by q-value < 0.01 for all group comparisons in this study. **Supplemental table 2** gives the unfiltered quantification results output from Spectronaut for the intracellular proteomics data. **Supplemental table 3** gives the intracellular proteomics raw data used by spectronaut to determine altered proteins, including peptide-level peak areas. **Supplemental table 4** gives the unfiltered quantification results output from Spectronaut for the secretome proteomics data. **Supplemental table 5** gives the secretome proteomics raw data used by spectronaut to determine altered proteins, including peptide-level peak areas.

---

**Supplementary Figure 1: RPE intracellular proteome changes resulting from chronic ROS (3 weeks PQ treatment). (**A) Barplots showing quantification of the 6 most increased proteins. (B) Barplots showing quantification of the 6 most decreased proteins. Error bars show the standard error. Significant changes defined as at least 1.5-fold and q-value<0.05 (*), q-value < 0.01 (**), and q-value<0.001 (***).

**Supplementary Figure 2: Notable secreted protein changes. (**A) Transforming growth factor β1 (TGF-β1), transforming growth factor β2 (TGF-β2), and bone morphogenic protein 1 (BMP1). (B) Apolipoprotein E (APOE), amyloid β-like protein 2 (APLP2), and urotensin-2 (UTS2). (C) Secreted proteases MMP2, ADAM9, and SCPEP1. Error bars show the standard error. Significant changes defined as at least 1.5-fold and q-value<0.05 (*), q-value < 0.01 (**), and q-value<0.001 (***).

