## Supplemental Figures for "Proteome and Secretome Dynamics of Stem Cell-Derived Retinal Pigmented Epithelium in Response to Acute and Chronic ROS"

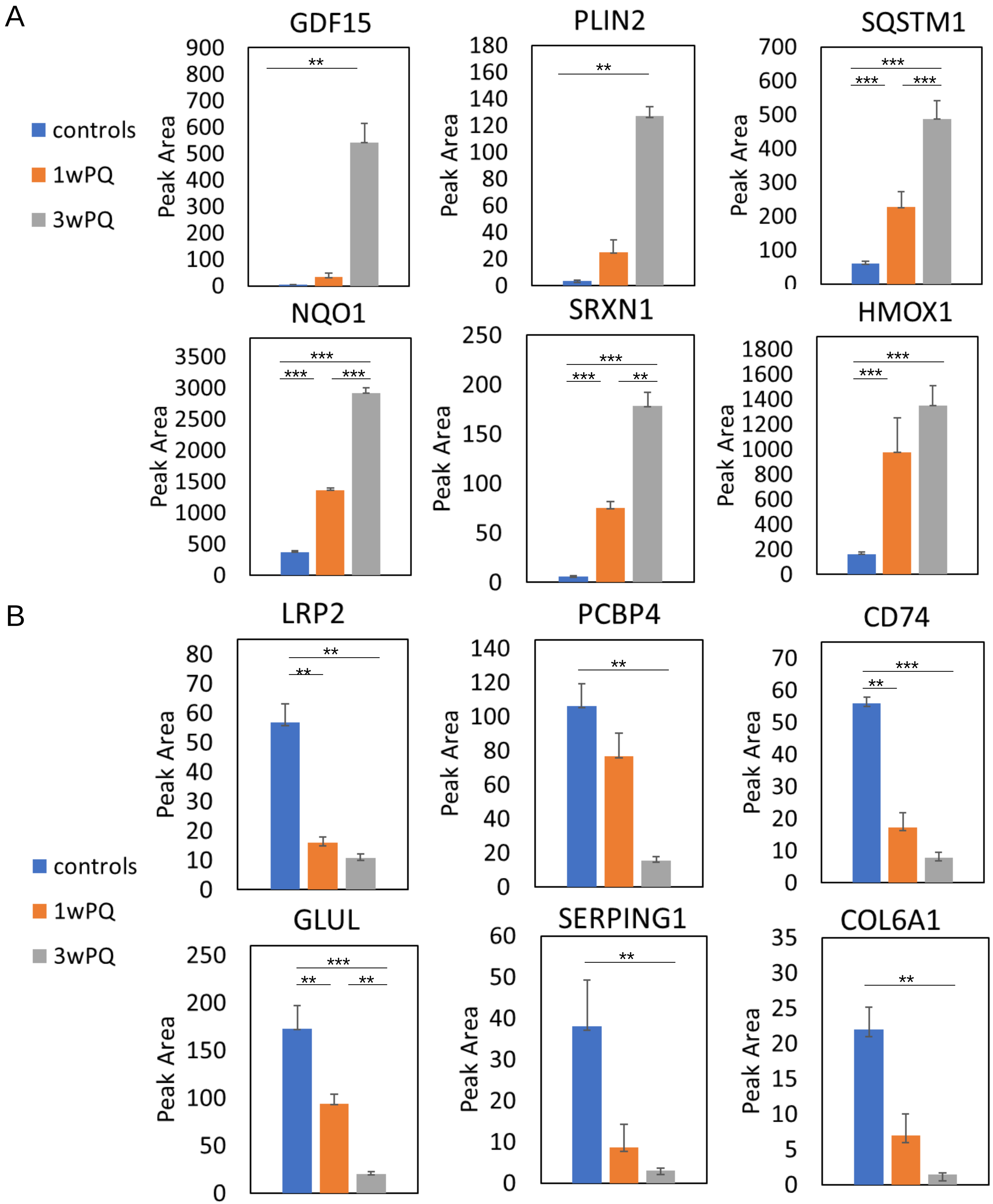


**Supplementary Figure 1: RPE intracellular proteome changes resulting from chronic ROS (3 weeks PQ treatment).** (a) Barplots showing quantification of the 6 most increased proteins. (b) Barplots showing quantification of the 6 most decreased proteins. Error bars show the standard error. Significant changes defined as at least 1.5-fold and q-value<0.05 (*), q-value < 0.01 (**), and q-value<0.001 (***).

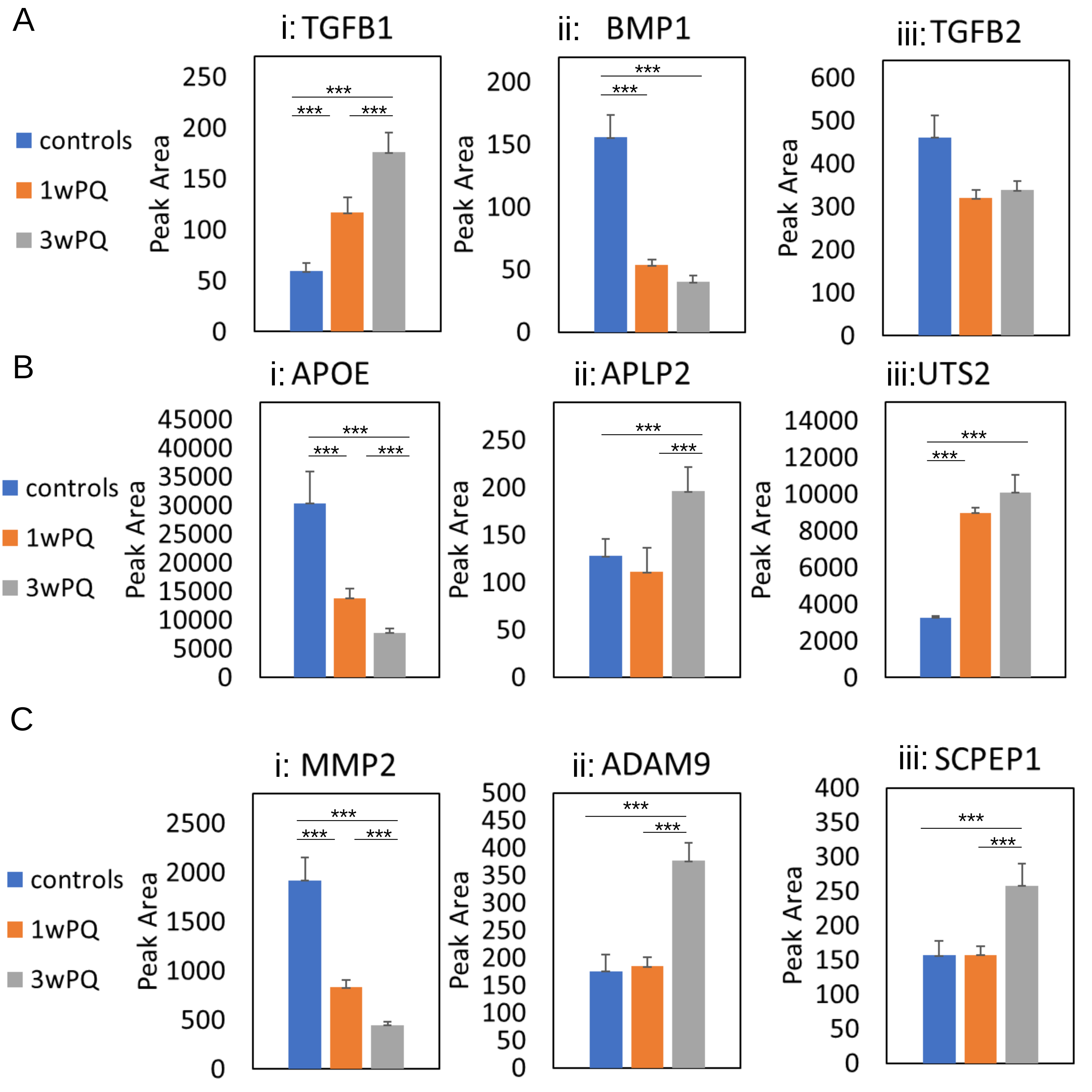


**Supplementary Figure 2: Notable secreted protein changes.** (A) Transforming growth factor β1 (TGF-β1), transforming growth factor β2 (TGF-β2), and bone morphogenic protein 1 (BMP1). (B) Apolipoprotein E (APOE), amyloid β-like protein 2 (APLP2), and urotensin-2 (UTS2). (C) Secreted proteases MMP2, ADAM9, and SCPEP1. Error bars show the standard error. Significant changes defined as at least 1.5-fold and q-value<0.05 (*), q-value < 0.01 (**), and q-value<0.001 (***).
